# Endothelial NCK2 Promotes Atherosclerosis Progression in Male but not Female *Nck1*-null Atheroprone Mice

**DOI:** 10.1101/2022.06.30.498326

**Authors:** Briana C. Bywaters, Gladys Pedraza, Andreea Trache, Gonzalo M. Rivera

## Abstract

A better understanding of endothelial dysfunction holds promise for more effective interventions for atherosclerosis prevention and treatment. Endothelial signaling by the non-catalytic region of the tyrosine kinase (NCK) family of adaptors, consisting of NCK1 and NCK2, has been implicated in cardiovascular development and postnatal angiogenesis but its role in vascular disease remains incompletely understood. Here, we report stage- and sex-dependent effects of endothelial NCK2 signaling on arterial wall inflammation and atherosclerosis development. Male and female *Nck1*-null atheroprone mice enabling inducible, endothelial-specific *Nck2* inactivation were fed a high fat diet (HFD) for 8 or 16 weeks to model atherosclerosis initiation and progression, respectively. Analysis of aorta preparations *en face* during disease progression, but not initiation, showed a significant reduction in plaque burden in males, but not females, lacking endothelial NCK2 relative to controls. Markers of vascular inflammation were reduced by endothelial NCK2 deficiency in both males and females during atherosclerosis progression but not initiation. At advanced stages of disease, plaque size and severity of atherosclerotic lesions were reduced by abrogation of endothelial NCK2 signaling only in males. Collectively, our results demonstrate stage- and sex-dependent modulation of atherosclerosis development by endothelial NCK2 signaling.

## Introduction

Endothelial dysfunction is a critical initiating event in the pathogenesis of atherosclerosis. It is well-established that atherosclerosis risk factors, in particular altered metabolism (1), promote a proinflammatory endothelium in regions of the arterial tree exposed to disturbed hemodynamics (2). It is also known that loss of atheroprotective endothelial functions is linked to increased expression of cell adhesion and chemoattractant molecules that foster the recruitment of leukocytes and their migration into the subintimal space (3). However, the mechanisms linking these risk factors to atherogenesis remain incompletely understood. Current therapies for atherosclerosis, relying primarily on reducing circulating levels of cholesterol (4) and inflammation (5), do not preserve or restore endothelial homeostasis. Such limitations, together with a shifting focus from treatment to prevention of atherosclerotic cardiovascular disease, underscore the need for a deeper understanding of the molecular mechanisms underpinning endothelial dysfunction.

The non-catalytic region of the tyrosine kinase (NCK) family of adaptors, consisting of NCK1 and NCK2, links tyrosine phosphorylation with cytoskeletal remodeling [reviewed in (6)]. Their architecture consists of three N-terminal Src Homology (SH)3 domains and a C-terminal SH2 domain tethered by unstructured linkers. NCK adaptors assemble multimolecular signaling complexes to promote endothelial cell migration (7) and morphogenesis (8) *in vitro*. Endothelial NCK signaling is required for cardiovascular development (9), postnatal retinal angiogenesis and pathological ocular neovascularization (10). Furthermore, NCK1 and NCK2 were shown to mediate oxidative stress-induced endothelial inflammation (11). Functional redundancy of NCK1 and NCK2 has been inferred by their conserved (∼68% identity) amino acid sequence (12) and absence of an overt phenotype in mice lacking either one of the paralogues (13). However, recent studies suggest that NCK1, but not NCK2, mediates disturbed flow-induced endothelial permeability (14), as well as atherogenic inflammation and plaque formation in male mice (14). To our knowledge, the role of endothelial NCK signaling in disease stage- and sex-dependent regulation of arterial wall inflammation and atherogenesis has not been investigated. In the present study, we show that inducible, endothelial-specific deletion of NCK2 does not impact HFD-induced atherosclerosis initiation. During atherosclerosis progression, however, abrogation of endothelial NCK2 signaling ameliorates inflammation of the arterial wall in both sexes, whereas severity of lesions and atherosclerosis progression are reduced in male, but not female, *Nck1*-null atheroprone mice.

## Materials and Methods

Information about antibodies and reagents is detailed in *Supplementary Materials and Methods*.

### Mice

Mice were maintained in the Comparative Medicine Program Laboratory Animal Resources and Research facility at Texas A&M University. Animal handling and experimental procedures were performed according to a protocol (AUP 2019-0184) approved by the Institutional Animal Care and Use Committee of Texas A&M University. *Nck1*^-/-^; *Nck2*^fl/fl^; *Cdh5*-Cre^ERT2^ mice were a kind gift from Anne Eichmann. These animals were generated by breeding *Nck1*^-/-^; *Nck2*^fl/fl^ mice (15, 16) with the *Cdh5*-Cre^ERT2^ line (17). We crossed *Nck1*^-/-^; *Nck2*^fl/fl^; *Cdh5*-Cre^ERT2^ mice with *ApoE*^-/-^ mice, purchased from The Jackson Laboratory (Bar Harbor, Maine), to generate mixed background *Nck1*^-/-^; *Nck2*^fl/fl^; ApoE^-/-^ mice with (eNck2-) and without (eNck2+) the *Cdh5*-Cre^ERT2^ driver. The Cre allele was passed through male breeders to prevent recombination in oocytes. DNA from tail snips was used for genotyping. At 8 weeks of age, all mice received a total of five intraperitoneal injections, administered every other day, of 75 mg/kg tamoxifen (Sigma-Aldrich) in 100 μL corn oil. After a one-week recovery period, all mice were subjected to a HFD, containing 0.2% cholesterol and 21% total fat (Envigo TD.88137), and water *ad libitum*. Mice were maintained on a 12-hour light/dark cycle. Both male and female animals were used in our study.

### Endothelial Cell Isolation and Western Blotting

Successful *Nck2* recombination was confirmed by detection of NCK protein levels in extracts from lung endothelial cells isolated as previously described (10). Briefly, lung tissue from two mice per genotype (eNck2+ and eNck2-) was enzymatically digested and endothelial cells were enriched with anti-CD31 coated Dynabeads. A second enrichment with anti-CD102-coated Dynabeads was performed before culturing cells at 37Cº on fibronectin-coated tissue culture dishes in EGM-2 media (Lonza Walkersville). Cells (passage two) were lysed in kinase lysis buffer and mixed with 5X SDS loading buffer. The mixtures were boiled at 95C for 5 minutes. Proteins were separated by 12% SDS-PAGE and transferred to a nitrocellulose membrane for western blotting using antibodies against CD144, NCK, and β-actin.

### Blood Serum Assays

Mice were fasted for 16 hours before blood sample collection via cardiac puncture into untreated tubes immediately following CO2 asphyxiation. Commercially available kits were used to determine serum levels of cholesterol (Invitrogen) and triglycerides (Invitrogen) following manufacturer’s instructions.

### Tissue Processing

To investigate atherosclerotic lesion properties, mice were perfused with ice cold DPBS followed by 10% non-buffered formalin (NBF) through the left ventricle. The heart and thoracic aorta were collected and kept in 10% NBF at 4Cº for 24 hours. We chose to analyze the aortic root to detect incipient atherosclerotic lesions because lesion development tends to occur earlier at this site compared to other sites of the arterial tree (18). The base of the heart was separated, and paraffin embedded 5 μm serial sections of the aortic root were collected for comparative analysis, as described previously (18). Slides were stained with H&E, Picrosirius Red, or used for fluorescence staining. Slides subjected to histologic staining were imaged in brightfield using an Olympus VS120 Virtual Slide Scanning System equipped with a U Plan S-Apo dry 10X objective. Fluorescence staining was performed using Isolectin-B4 and antibodies against ICAM-1, CD31, VCAM-1, and Mac-2. Preparations subjected to fluorescence staining were imaged using an Olympus Fluoview FV3000 Confocal Laser Scanning Microscope equipped with a U Plan S-Apo dry10X objective.

### Atherosclerotic Lesion Assessment

Aortas were cleaned of adventitial fat and stained with Oil Red O (Fisher). Vessels were then opened *en face* and imaged (3X) using a 16-megapixel LG camera. Analysis of *en face* images from aortas was performed using ImageJ software. Analysis of aortic root sections was completed using QuPath software. Samples that did not display all three leaflets of the aortic valve across three serial slides were excluded from analysis. Lesion area was determined by subtracting the area of the lumen from the area delimited by the internal elastic lamina. The sum of lesion area across serial sections with visible valve leaflets was calculated. All other quantifications were performed using a single representative slide containing four serial sections for each sample. For quantification of collagen, Picrosirius Red signal was detected using a standardized pixel classification thresholder. Medial area was calculated as the difference between the areas delimited by the internal and external elastic laminae. Lesions were classified according to established criteria (19). For fluorescence images, a pixel classifier thresholder was developed for each marker and applied to all corresponding samples. For samples stained for CD31 and ICAM-1 as well as VCAM-1 and Mac-2, the neointima was traced by hand using the internal elastic lamina as the outer bound.

### Statistical Analyses

Normal distribution assumptions were assessed using the Shapiro Test. Datasets that followed a normal distribution were subjected to two-way ANOVA followed by Tukey’s multiple means comparison test. Datasets that did not follow a normal distribution were analyzed using Wilcoxon Rank Sum Test with Bonferroni correction. Differences were considered significant at *p* <0.05. Statistical analysis and visualization were performed using R v4.0.5. Quantitative results are displayed using box plots, where the bottom and top horizontal lines in boxes indicate the 25^th^ and 75^th^ percentile, respectively, and the central line is the median. Whiskers extend 1.5 x the interquartile range and dots represent individual observations. Numbers of animals/group (n) are indicated in parentheses.

## Results

### Endothelial NCK2 accelerates atherosclerosis progression in male mice

We crossed APOE-deficient mice with mice homozygous for both an *Nck1*-null allele and a floxed *Nck2* allele, and that carried tamoxifen-inducible Cre recombinase expressed under control of the *Cdh5* promoter (Fig. 1A). In these animals, tamoxifen-induced Cre activity results in *Nck2* inactivation by excision of the loxP site-flanked exon 1 (Fig 1B). The resulting progeny of mixed genetic background consisted of atheroprone mice either lacking (*Nck1*^*-/-*^; *Nck2*^*fl/fl*^; *Apoe*^*-/-*^) or carrying (*Cdh5*-Cre^ERT2^; *Nck1*^*-/-*^; *Nck2*^*fl/fl*^; *Apoe*^*-/-*^) Cre recombinase, herein termed eNck2+ and eNck2- (Fig. 2C, left), respectively. Tamoxifen injection almost completely abolished NCK2 expression selectively in the endothelium of eNck2-but not eNck2+ animals (Fig 1C, right). To ascertain specific roles of NCK2 signaling and potential disease stage- and sex-dependent differences in endothelial inflammation and atherosclerosis, eight-week-old male and female mice from both genotypes were treated with tamoxifen and subsequently fed a HFD for 8 or 16 weeks to model atherosclerosis initiation and progression, respectively (Fig. 1D). Serum triglyceride and cholesterol levels did not differ by genotype or sex after 8 or 16 weeks of HFD (Suppl. Fig. 1). Similarly, body weight gain did not differ by genotype or sex after 8 weeks of HFD. Regardless of genotype, however, males gained more weight than females after 16 weeks of HFD (Suppl. Fig. 1). Plaque burden, determined in *en face* preparations of thoracic aortas stained with Oil Red O (Fig. 1E), did not differ by genotype or sex after 8 weeks of HFD (Fig. 1F). Although the overall plaque burden increased, only male mice lacking endothelial NCK2 (eNck2-) displayed decreased (*p*<0.05) plaque burden compared to control eNck2+ animals after 16 weeks of HFD (Fig. 1F). Thus, these results suggest an important, sex-dependent role for endothelial NCK2 in atherosclerosis progression but not initiation.

**Figure 1.**
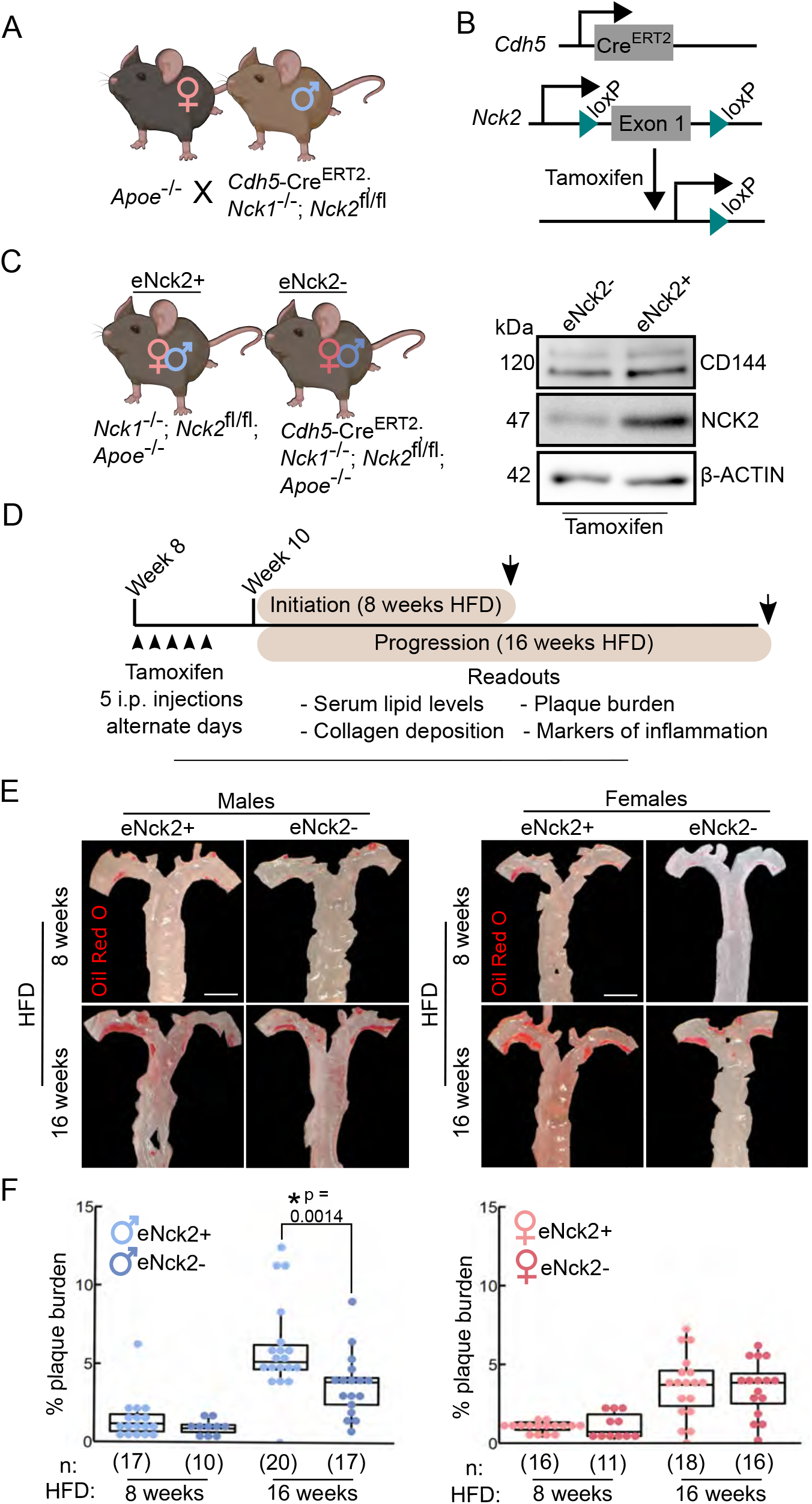
Assessment of atherosclerosis initiation and progression in atheroprone male and female mice with or without endothelial-specific deletion of NCK2. **(A)** APOE-deficient (*Apoe*^-/-^) mice were crossed with mice homozygous for both an *Nck1*-null allele and a floxed *Nck2* allele and that carried inducible Cre recombinase to selectively inactivate *Nck2* in the endothelium (*Cdh5-Cre*^ERT2^; *Nck1*^-/-^; *Nck2*^fl/fl^). **(B)** Tamoxifen-induced Cre recombination mediates excision of the loxP site-flanked exon 1 of *Nck2*. **(C)** Experimental groups consisted of male and female mice with (eNck2+) and without (eNck2-) NCK2 expression in the endothelium (left). Western blots using extracts from lung endothelial cells showing tamoxifen-induced deletion of NCK2 alongside levels of the endothelial marker CD144 (VE-cadherin) and the loading control β-ACTIN (right). **(D)** Diagram showing timeline (starting on week 8 after birth) of experimental manipulations, including tamoxifen injections (arrowheads), duration of high fat diet (HFD) regime, timing of tissue collection (arrows), and key readouts. **(E)** Representative images of *en face* preparations of mouse thoracic aortas stained with Oil Red O. Aortas were obtained from male (left) and female (right) mice fed HFD for 8 or 16 weeks to assess atherosclerosis initiation and progression, respectively. The scale bar represents 2 mm. **(F)** Box plots showing quantification of plaque burden as percentage of exposed intimal area occupied by lesions in male (left) and female (right) mice. Statistical significance was determined by two-way ANOVA followed by Tukey’s multiple means comparison test. \**p* < 0.05. Numbers of animals/group (n) are indicated in parentheses.

**Figure 2.**
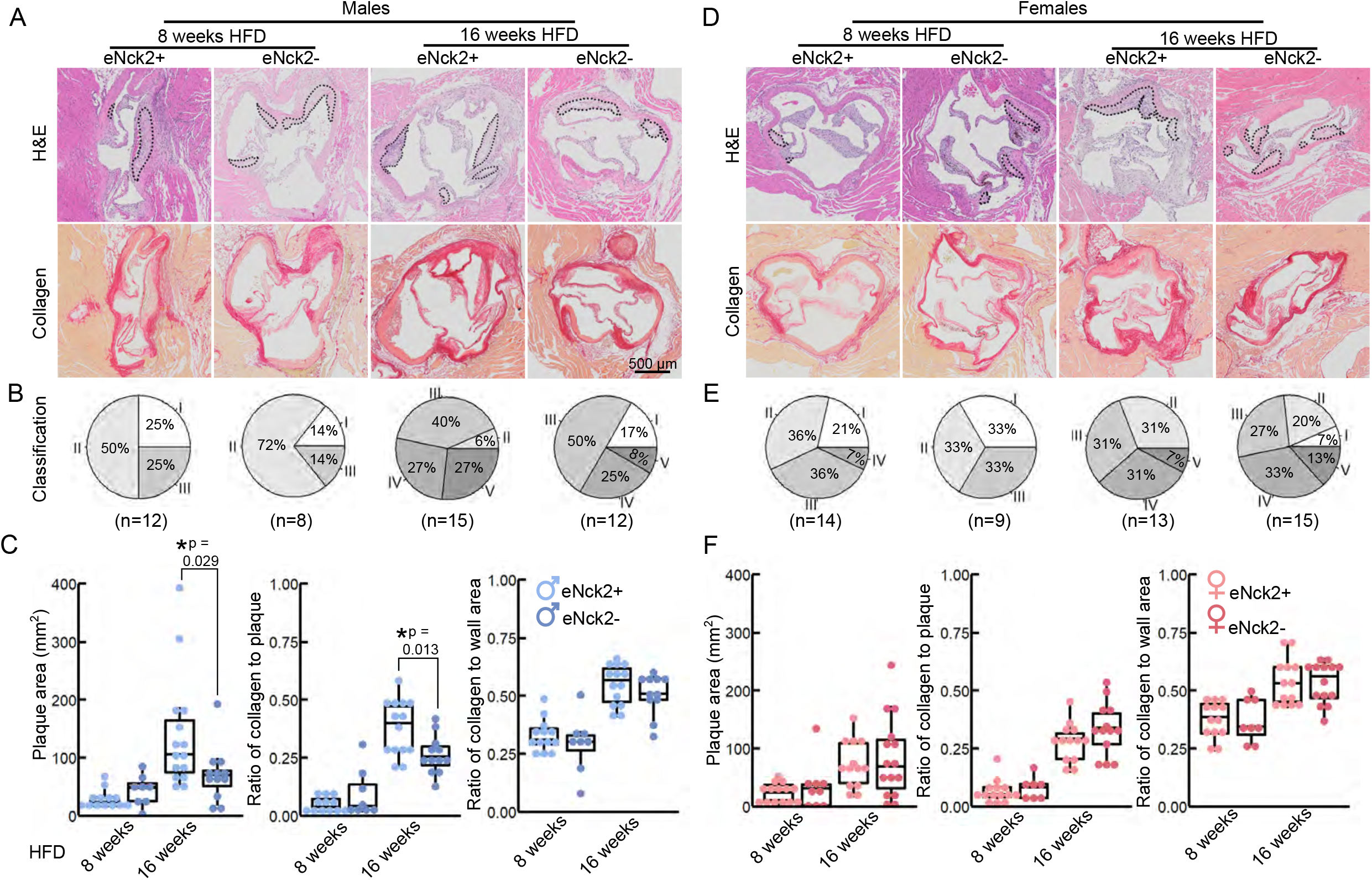
Morphometric analysis of aortic root lesions in atheroprone male and female mice with and without endothelial-specific deletion of NCK2. Animals were fed a HFD for 8 or 16 weeks to assess atherosclerotic lesion initiation and progression, respectively. **(A**/**D)** Representative images of aortic roots stained with H&E (top) and Picrosirius Red (bottom) to assess lesion development and collagen deposition, respectively. The scale bar represents 500 μm. **(B/E)** Pie charts showing percentage of lesion category in each experimental group. Lesions were classified as intima thickening (type I), fatty streaks (type II), intermediate lesion (type III), advanced atheroma (type IV), and fibroatheroma (type V). Numbers of animals/group are indicated in parentheses. **(C/F)** Box plots showing quantification of plaque area, ratio of collagen to plaque area, and ratio of collagen to aortic root wall area. Statistical significance was determined by Wilcoxon Rank Sum Test with Bonferroni correction. \**p* < 0.05.

### Endothelial NCK2 increases plaque development and lesion severity in male mice

We used H&E and Picrosirius Red staining to analyze atherosclerotic lesion development and collagen deposition in aortic roots of male and female mice fed HFD for 8 or 16 weeks. Both lesion development and collagen deposition increased with prolonged exposure to HFD regardless of sex or genotype (Fig. 2A and D). For each experimental condition, defined by the duration of HFD feeding, genotype, and sex, we determine the percentage of lesion type based on progression of pathological changes as previously described (19): intima thickening (type I), fatty streaks (type II), intermediate lesion (type III), advanced atheroma (type IV), fibroatheroma (type V). A slightly slower progression of pathological changes at initial disease stages (8 weeks of HFD) was seen in endothelial NCK2-deficient male (Fig. 2B), but not female (Fig. 2E), mice. As expected, advanced pathological changes were associated with the prolonged (16 weeks) regime of HFD. However, complete absence of early stage (type I) alongside a higher percentage of late stage (type V) lesions was observed in male (Fig.2 B) but not female (Fig. 2E) mice expressing endothelial NCK2 (male eNck2+ vs. eNck2-, *p*<0.05). Furthermore, male eNck2+ mice developed significantly more plaque and accumulated more collagen in plaques than the eNck2-counterpart (Fig. 2C, left & center panels). A larger proportion of eNck2+ vs. eNck2-males also developed lesions with clearly identifiable necrotic core after 16 weeks of HFD (Suppl. Fig. 2). The incidence of other pathological findings, including media erosion and aortic dissection, did not differ by disease stage, sex, or genotype (Suppl. Fig. 2). Although the ratio of collagen to wall area increased with the prolonged HFD regime in males, no genotype-dependent changes were observed (Fig. 2C, right panel). In females, increase in plaque area, collagen content in plaques, and the ratio of collagen to wall area were dependent on the duration of the HFD regime but independent of genotype (Fig. 2F). Consistent with the assessment of plaque burden (Fig. 1E-F), analysis by histomorphometry shows that endothelial NCK2 promotes plaque development, i.e., increased plaque size and advancement of pathological changes, in male but not female atheroprone mice fed HFD for 16 weeks.

### Inactivation of endothelial NCK2 reduces expression of ICAM-1 in advanced disease stage

We performed co-staining of the endothelial and cell adhesion marker, CD31 and ICAM-1 respectively, in aortic roots obtained from male and female mice from both genotypes fed HFD for 8 or 16 weeks (Fig. 3A and C). Total ICAM-1 expression, defined as the area of ICAM-1 signal in the entire aortic root normalized to the aortic root area, did not differ by genotype or sex at initial stages (8 weeks HFD) of disease (Fig. 3B and D, left panels). In animals fed HFD for 16 weeks, however, inactivation of endothelial NCK2 decreased (eNck2+ vs. eNck2-, *p*<0.05) total ICAM-1 expression in both males and females (Fig. 3B and D, left panels). We assessed neointimal ICAM-1 expression, which was defined as the area of the ICAM-1 signal in the neointima normalized by the neointima area, which is delimited by the internal elastic lamina. At early stages of disease (8 weeks HFD), inactivation of endothelial NCK2 decreased neointimal ICAM-1 expression in males, but not females. At advanced stages of disease (16 weeks of HFD), inactivation of endothelial NCK2 decreased ICAM-1 neointimal expression in both males and females (eNck2+ vs. eNck2-, *p*<0.05). Because endothelial cell ICAM-1 surface expression is elevated at sites of endothelial activation (20), we also determined intimal ICAM expression, which was defined as the area of the ICAM-1 signal normalized to the area of CD31 signal in a 10 μm-thick periluminal layer (intima). Intimal ICAM-1 expression did not differ by genotype or sex in animals fed HFD for 8 weeks. At advanced stages of disease (16 weeks of HFD), inactivation of endothelial NCK2 decreased ICAM-1 intimal expression in both males and females (eNck2+ vs. eNck2-, *p*<0.05). In sum, endothelial NCK2 signaling promotes atherosclerosis progression, but not initiation, through a mechanism that involves increased expression of ICAM-1 in the arterial wall and neointimal/intimal layers.

**Figure 3.**
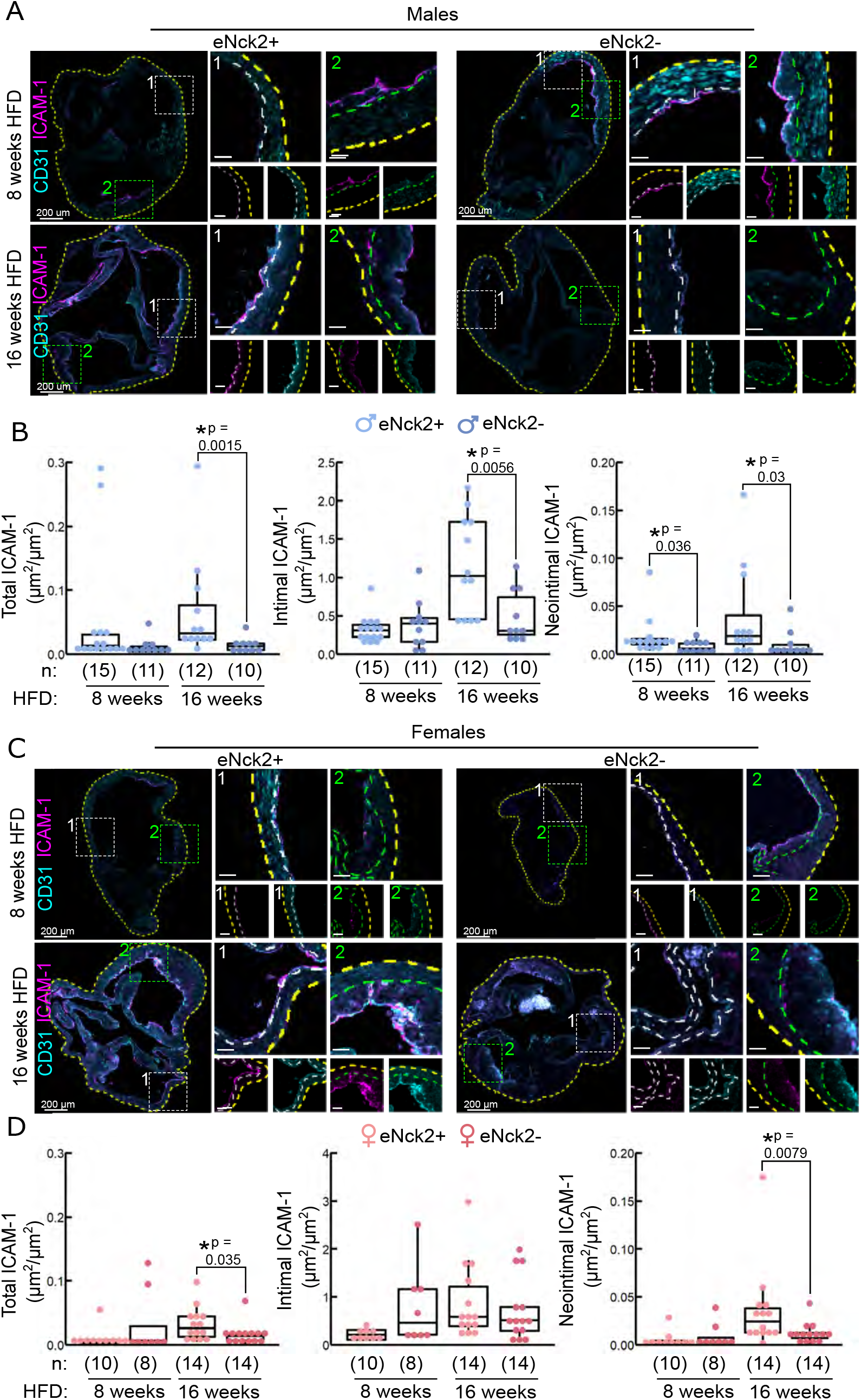
Determination of ICAM-1 in aortic roots of atheroprone mice with and without endothelial-specific deletion of NCK2. Animals were fed a HFD for 8 or 16 weeks to assess atherosclerotic lesion initiation and progression, respectively. **(A/C)** Representative confocal images of aortic roots showing immunofluorescence staining of endothelial (CD31) and cell adhesion (ICAM-1) markers in males **(A)** and females **(C)**. The dotted yellow, white, and green lines contour the aortic root section, intimal, and neointima layers, respectively. The scale bars represent 50 μm unless otherwise indicated. **(B/D)** Box plots showing quantification of total (left panels), intimal (center panels), and neointimal (right panels) ICAM-1 in males **(B)** and females **(D)**. Total ICAM-1 was calculated as ICAM-1 area in entire root/root area. Intimal ICAM-1 was calculated as the area of ICAM-1/area of CD31 in signal in a 10 μm-thick periluminal layer (intima). Neointimal ICAM-1 was calculated as area of ICAM-1/neointimal area. Dots represent individual animals and the number of animals per group is indicated in parentheses. Statistical significance was determined by Wilcoxon Rank Sum Test with Bonferroni correction. \**p* < 0.05.

### Endothelial Nck2 promotes vascular wall inflammation

To further assess inflammatory changes in the arterial wall, we co-stained VCAM-1 (inflammation marker), Mac-2 (macrophage infiltration), and the endothelium (IB4) in aortic roots obtained from male (Fig. 4A) and female (Fig. 4C) mice from both genotypes fed HFD for 8 or 16 weeks. We quantified total VCAM-1 expression (VCAM-1) and macrophage infiltration (Mac-2), respectively, as the area of VCAM-1 or Mac-2 signal in the entire root normalized to the root area. In addition, we quantified neointimal expression of VCAM-1 and macrophage infiltration as the area of VCAM-1 or Mac-2 signal in the neointima normalized to the neointimal area, which is delimited by the internal elastic lamina. Although enriched in atherosclerotic lesions, total (Fig. 4B and D, bottom left panels) and neointimal (Fig. 4B and D, bottom right panels) VCAM-1 expression did not differ by genotype, sex, or duration of HFD regime. Similarly, total and neointimal macrophage infiltration, assessed by Mac-2 staining, did not differ by genotype or sex in animals fed HFD for 8 weeks (Fig. 4B and D). In contrast, in animals fed HFD for 16 weeks, total (Fig. 4B and D, top left panels) and neointimal (Fig. 4B and D, right panels) macrophage infiltration was significantly higher in both male and female control mice compared to those lacking endothelial NCK2 (eNck2+ vs. eNck2-, *p*<0.05). Thus, endothelial NCK2 signaling exacerbates macrophage infiltration during advanced stages of atherosclerotic disease.

**Figure 4.**
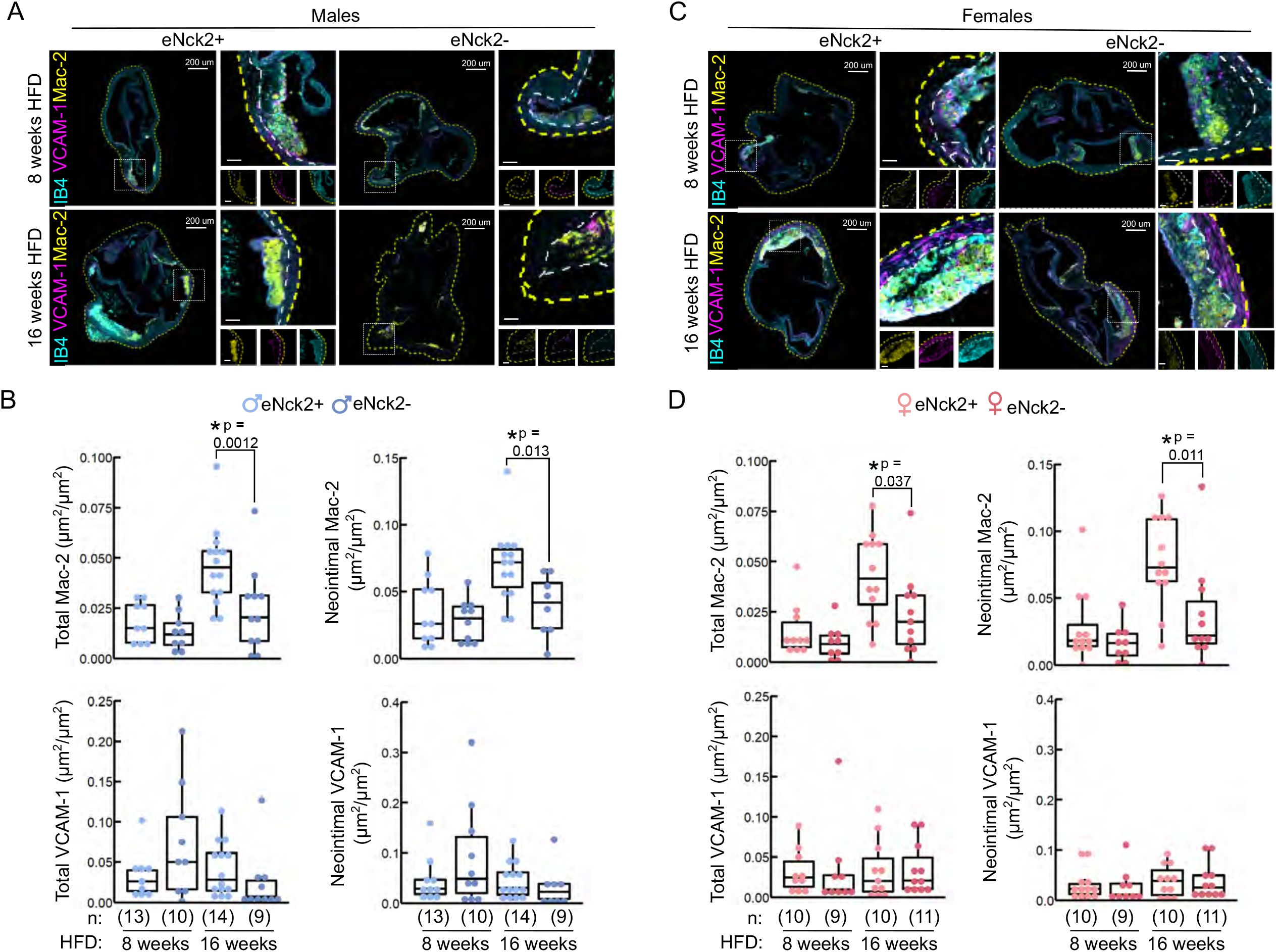
Inflammation in aortic roots obtained from atheroprone mice with or without endothelial-specific deletion of NCK2. Animals were fed a HFD for 8 or 16 weeks to assess atherosclerotic lesion initiation and progression, respectively. **(A/C)** Representative confocal images of aortic roots showing immunofluorescence staining of the cell adhesion (VCAM-1) and leukocyte infiltration (Mac-2) markers in males **(A)** and females **(C)**. The dotted yellow and white lines contour the aortic root section and the neointima, respectively. The scale bars represent 50 μm unless otherwise indicated. **(B/D)** Box plots displaying quantification of total Mac-2 or VCAM (left panels) and neointimal Mac-2 or VCAM-1 (right panels) in males **(B)** and females **(D)**. Total Mac-2 and VCAM-1 were calculated as Mac-2 or VCAM-1 area in entire root/root area. Neointimal Mac-2 and VCAM-1 were calculated as Mac-2 or VCAM-1 area in the neointima/area of neointima. Dots represent individual animals and the number of animals per group is indicated in parentheses. Statistical significance was determined by Wilcoxon Rank Sum Test with Bonferroni correction. \**p* < 0.05.

## Discussion and Conclusions

Targeting mechanisms underlying endothelial activation and dysfunction holds promise for the development of novel and more effective interventions for the prevention and treatment of atherosclerosis. Implementing a robust experimental design enabling analysis of atherosclerosis initiation and progression in both male and female *Nck1*-null atheroprone mice, we show here that endothelial NCK2 exerts disease stage- and sex-dependent effects on arterial inflammation and atherosclerosis development. Specifically in males, NCK2 signaling in the endothelium promotes HFD-stimulated vascular wall inflammation and atherosclerotic progression but not initiation. In females, however, these effects appear to be decoupled. Although also dispensable for atherosclerosis initiation, in advanced disease stages endothelial NCK2 signaling in females contributes to HFD-induced vascular wall inflammation but not plaque development. These results point to the increasingly recognized significance of sexual dimorphism and temporal regulation as critical factors underlying the complexities of atherosclerosis pathogenesis.

The highly similar NCK1 and NCK2 adaptors, sharing 68% amino acid identity, have overlapping functions during development (13). Expression of NCK2 in the endothelium compensates the global inactivation of *Nck1* during cardiovascular development (9) and postnatal angiogenesis (10). Our detailed analysis of *Nck1*-null atheroprone mice shows a significant role of endothelial NCK2 signaling in regulation of arterial wall inflammation, as evidenced by increased ICAM-1 expression and macrophage infiltration but not VCAM-1 expression. The absence of genotype-dependent differences in VCAM-1 expression in our study, is consistent with the highly variable expression of VCAM-1 in both intact areas and atherosclerotic lesions of the aortic root previously described in APOE-deficient mice fed HFD (21). Interestingly, despite being present in both sexes, decreased arterial wall inflammation translated into reduced plaque burden (en face aortas), plaque area and severity of atherosclerotic lesions (aortic roots) only in males lacking endothelial NCK2. Although the mechanism underlying the sex-specific differences in atherosclerosis progression described in this study remains to be elucidated, sexual dimorphism in NCK signaling appears to be an emerging theme (22).

Strengths of the present study include analysis of early and advanced stages of HFD-induced atherosclerosis and direct statistical comparison of readouts between sexes. Our results, however, appear at variance with those of recent studies examining the role of NCK signaling in atherosclerosis. Implementing the partial carotid ligation model of endothelial dysfunction and atherosclerosis (23), Orr and colleagues recently showed that endothelial NCK1, but not NCK2, mediates atheroprone flow-induced endothelial dysfunction/increased permeability (24). In a follow up study based on analysis of male mice, the same laboratory proposed a specific role for NCK1, but not NCK2, in atheroprone flow- and HFD-induced atherogenic inflammation and plaque formation (14). Results from the latter study are, however, difficult to interpret considering that *i)* comparisons involved mice with different developmental levels of NCK, i.e., mice with constitutive, global inactivation of *Nck1* and mice with inducible, endothelial-specific inactivation of *Nck2* against controls expressing wild type levels of NCK1 and NCK2, *ii)* a decreased inflammatory response was coupled to global *Nck1* inactivation; specifically, plasma levels of several proinflammatory molecules, including interleukin 1α, interleukin 1β, tumor necrosis factor α, and C-C motif chemokine 2 (MCP-1), were reduced in mice with constitutive, global NCK1 deficiency but not in control mice or mice with endothelial-specific abrogation of NCK2 signaling, and *iii)* vascular inflammation and plaque burden were assessed at a single interval, 12 weeks of HFD, during disease progression We sought to establish a phenotype for plaque development in both males and females of the *Nck1*-null atheroprone mouse with or without inducible (post-developmental) endothelial-specific *Nck2* inactivation to allow for direct comparison. Thus, one limitation of the present study is that it does not explore the molecular underpinnings of the identified phenotype, which falls beyond of the scope of this brief research report. Full elucidation of the specific roles of NCK paralogues in vascular cell dysfunction and atherosclerosis pathogenesis warrants further investigation. To maximize impact, future studies should implement recommendations for the design and execution of preclinical studies of atherosclerosis to uncover mechanisms underlying initiation, propagation, and regression of atherosclerotic lesions (25), and enable well-powered, direct statistical comparison of sexes (26).

In conclusion, the present report identifies previously unappreciated roles of endothelial NCK2 signaling in stage- and sex-dependent modulation of atherosclerosis development.

## Data Availability Statement

The raw data originating the results and supporting the conclusions of this study are available upon reasonable request from the corresponding authors.

## Ethics Statement

All animal studies were reviewed and approved by the Texas A&M University Institutional Animal Care and Use Committee.

## Author Contributions

GR conceptualized the study. BB maintained the mouse colony, implemented the experimental design, and performed necropsies and tissue collection. BB and GP processed tissues. BB, AT and GR analyzed data and interpreted results. BB and GR prepared the manuscript. All authors approved the final version of the manuscript.

## Acknowledgments

We thank Dr. Anne Eichmann (Yale School of Medicine) for kindly providing the *Nck1*^-/-^; *Nck2*^fl/fl^; *Cdh5*-Cre^ERT2^ mice, Drs. Kyung Ae Ko and Jun-Ichi Abe (MD Anderson Cancer Center) for training in the handling of murine arterial tissues, and Ms. Denise Pedraza and Mr. Rizwan Mahmood for their assistance in data processing. The authors acknowledge the assistance of the College of Veterinary Medicine and Biomedical Sciences Core Histology Lab Research Unit (RRID:SCR_022201), Texas A&M University, and the Integrated Microscopy and Imaging Laboratory (RRID:SCR_021637), Texas A&M Health Science Center.

## Contribution to the Field Statement

Atherosclerosis remains the leading cause of death worldwide. Currently available therapeutics lower cholesterol and reduce inflammation. However, such therapies are unable to fully restore vascular integrity. These limitations underscore the need for a deeper understanding of factors and mechanisms leading to vascular disease. Dysfunction of the innermost layer of cells, the endothelium, is an independent risk factor for atherosclerosis. Unveiling mechanisms underlying endothelial dysfunction may provide novel prevention and therapeutic opportunities. Here, we assessed how inactivating a gene that contributes to regulation of the endothelium alters arterial inflammation and plaque development during atherosclerosis in male and female mice.

## Supplementary Materials

This report includes Supplementary Materials and two Supplementary Figures.

**Supplemental Figure 1**. Assessment of serum lipid profile and body weigh change in atheroprone male and female mice with or without endothelial-specific deletion of NCK2. Quantification of serum triglyceride levels (left), serum cholesterol levels (center) and weight gain (right) in animals fed HFD for 8 (top panels) or 16 (bottom panels) weeks. Number of mice per group is indicated in parentheses. Data presented as mean ± SD. Statistical significance was determined by two-way ANOVA with Tukey’s multiple comparison test. \**p* < 0.05

**Supplemental Figure 2**. Analysis of pathological findings in atheroprone male and female mice with or without endothelial-specific deletion of NCK2. Representative images (top panels) and percentage of mice/group (bottom panels) presenting plaques with necrotic cores **(A)**, medial erosion **(B)**, and aortic dissection **(C)**. The number of mice per group is indicated in parentheses. Statistical significance was determined by Fisher’s exact test. \**p* < 0.05.

