## Supplementary material for "Endothelial NCK2 Promotes Atherosclerosis Progression in Male but not Female *Nck1*-null Atheroprone Mice": Suppl.Figure1

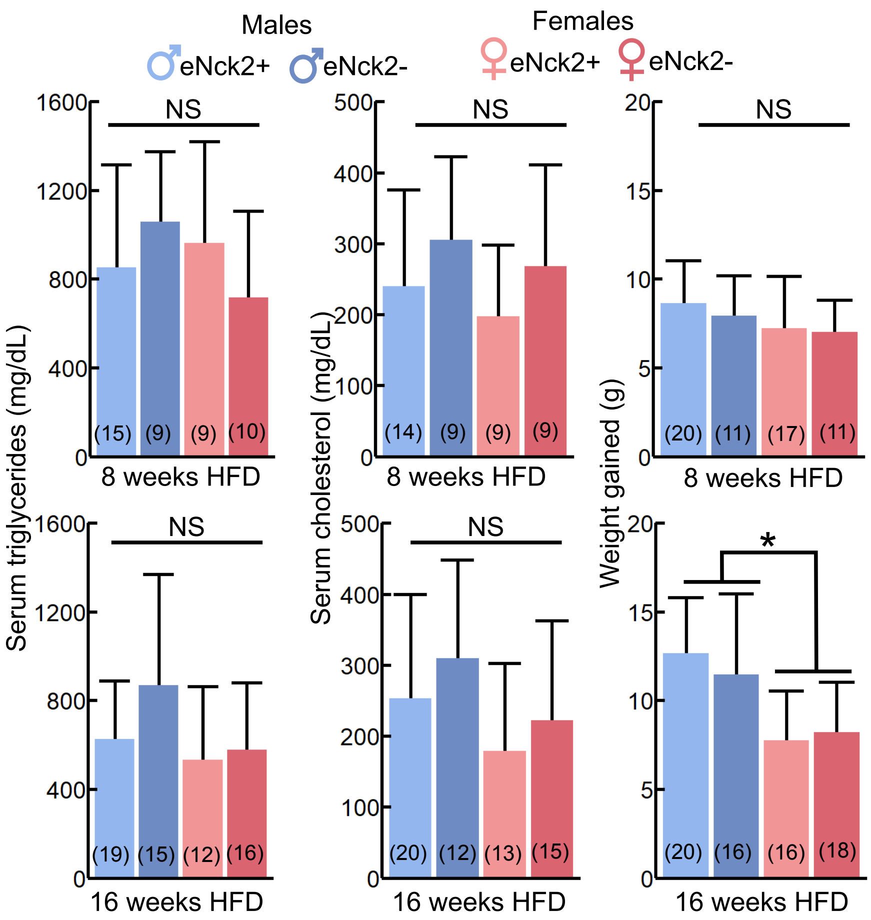


**Supplemental Figure 1.** Assessment of serum lipid profile and body weigh change in atheroprone male and female mice with or without endothelial-specific deletion of NCK2. Quantification of serum triglyceride levels (left), serum cholesterol levels (center) and weight gain (right) in animals fed HFD for 8 (top panels) or 16 (bottom panels) weeks. Number of mice per group is indicated in parentheses. Data presented as mean ± SD. Statistical significance was determined by two-way ANOVA with Tukey's multiple comparison test. *$p < 0.05$
