## Supplementary material for "Endothelial NCK2 Promotes Atherosclerosis Progression in Male but not Female *Nck1*-null Atheroprone Mice": Suppl.Figure2

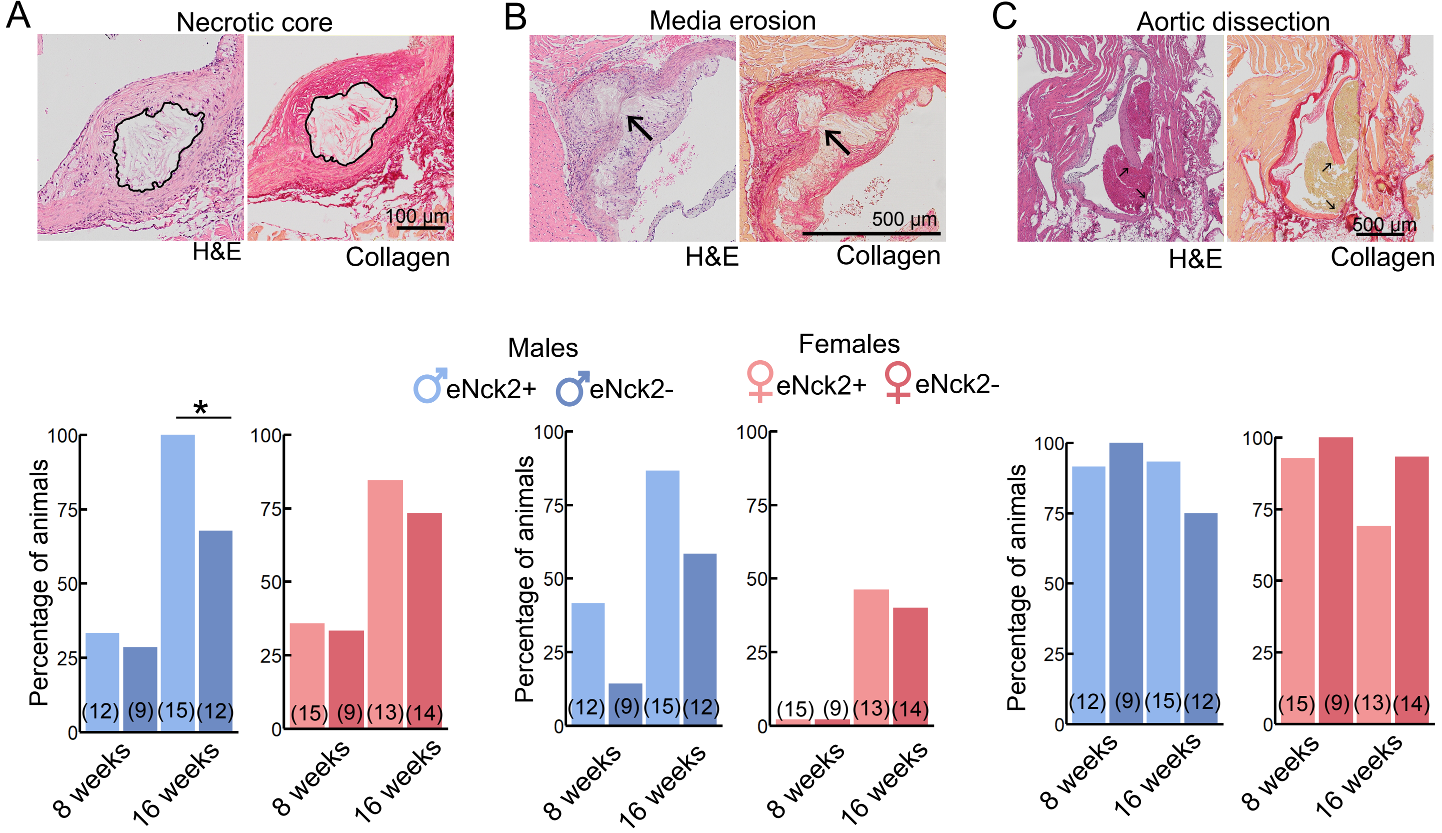


**Supplemental Figure 2**. Analysis of pathological findings in atheroprone male and female mice with or without endothelial-specific deletion of NCK2. Representative images (top panels) and percentage of mice/group (bottom panels) presenting plaques with necrotic cores **(A)**, medial erosion **(B)**, and aortic dissection **(C)**. Evidence of pathology is indicated in black outline **(A)** or with black arrows **(B-C)**. The number of mice per group is indicated in parentheses. Statistical significance was determined by Fisher's exact test. $^*p < 0.05$.
