## Supplemental Material for "Endothelial NCK2 Promotes Atherosclerosis Progression in Male but not Female *Nck1*-null Atheroprone Mice"

Supplementary Material

**PCR primers**

| **Gene** | **Forward** | **Reverse** |
| --- | --- | --- |
| Nck1 WT | GCATGTAGACAATTACACTTCAGCACC | ATTCATGGAATTTCGAACTCGCCACC |
| Nck1 mutant | CTGATTGAAGCAGAAGCCTGCGATG | TATTGGCTTCATCCACCACATACAGG |
| Nck2 WT | GAGGAATGCTGCCAACAGGACAGG | CACATACAGATACACACACGCTGAAG |
| Nck2 floxed | CTGATTGAAGCAGAAGCCTGCGATG | TATTGGCTTCATCCACCACATACAGG |
| ApoE | GCCTAGCCGAGGGAGAGCCG | WT: TGTGACTTGGGAGCTCTGCAGC Mutant: GCCGCCCCGACTGCATCT |
| Cre | TCCTGATGGTGCCTATCCTC | CCTGTTTTGCACGTTCACCG |

**Antibodies – Lung Endothelial Cell Isolation**

| **Antibody** | **Clonal** | **Species** | **Supplier** | **Catalog #** | **Concentration** | **Dilution** |
| --- | --- | --- | --- | --- | --- | --- |
| CD31 | Mono | Rat | BD Bioscience | 550274 | 15.625 µg/ml | 5 µg/mL |
| CD102 | Mono | Rat | BD Bioscience | 553326 | 0.5 mg/ml | 5 µg/mL |

**Other Reagents - Lung Endothelial Cell Isolation**

| **Reagent** | **Supplier** | **Catalog #** |
| --- | --- | --- |
| Collagenase/Dispase | Sigma Aldrich | 10269638001 |
| Dynabeads Protein G | Invitrogen | 10003D |
| Fibronectin - bovine plasma | R&D Systems | 1030-FN |
| EGM-2 cell culture media | Lonza Walkersville | CC-3162 |

**Buffers – Cell Lysis**

Kinase lysis buffer: 25mM pH 7.4 Tris, 150mM NaCl, 5mM EDTA, 1% Triton X-100, 10mM β-GP(β-glycerophosphate), 10mM Na3VO4, 10% Glycerol
- 1:500 0.5M phenylmethylsulfonyl fluoride, 1:100 Aprotinin, 1:500 50mM Na3VO4 added on use

5X SDS loading buffer: 5% β-Mercaptoethanol, 0.02% Bromophenol blue, 30% glycerol, 10% Sodium dodecyl sulfate in 250mM pH 6.8 Tris-Cl

**Antibodies – Western Blotting**

| **Antibody** | **Size** | **Clonal** | **Species** | **Supplier** | **Cat #** | **Dilution** | **Buffer** | **Duration** | **T°** |
| --- | --- | --- | --- | --- | --- | --- | --- | --- | --- |
| CD144 | 130 kDa | Mono | Mouse | Santa Cruz | sc-9989 | 1:1,000 | 3% NFDM | O/N | 4C |
| Nck | 47 kDa | Mono | Mouse | BD Bioscience | 610100 | 1:3,000 | 3% NFDM | O/N | 4C |
| B-actin | 42 kDa | Mono | Mouse | Sigma Aldrich | A5316 | 1:10,000 | 3% NFDM | O/N | 4C |
| IgG-HRP | -- | Poly | Goat | Santa Cruz | sc-2005 | 1:10,000 | 5% NFDM | 1 hour | RT |
| IgG-HRP | -- | Poly | Goat | Santa Cruz | sc-2004 | 1:10,000 | 5% NFDM | 1 hour | RT |

*NFDM:* Non-fat dry milk in TBS-T

**Other Reagents – Western Blotting**

| **Reagent** | **Supplier** | **Catalog #** |
| --- | --- | --- |
| 30% Acrylamide Solution | Protogel | EC890 |
| TEMED | Thermo Scientific | 17919 |
| Aprotinin | Sigma Aldrich | A6279 |
| BioTrace NT Nitrocellulose | Pall | 66485 |
| Western Lightning Plus chemiluminescent substrate | Perkin Elmer | NEL103E001EA |

**Reagents - Mice**

| **Reagent** | **Supplier** | **Catalog #** |
| --- | --- | --- |
| Tamoxifen | Sigma Aldrich | T5648 |
| HFD (15.2% protein, 42.7% CHO, **42% fat**) | Envigo | TD.88137 |

**Reagents - Lipid Analysis**

| **Reagent** | **Supplier** | **Catalog #** |
| --- | --- | --- |
| Amplex Red Cholesterol Assay Kit | Invitrogen | A12216 |
| Infinity Triglycerides Liquid Stable Reagent | Invitrogen | TR22421 |

**Antibodies - Immunohistochemistry**

| **Antibody** | **Clonal** | **Species** | **Supplier** | **Catalog #** | **Dilution** | **Duration** | **Temp** |
| --- | --- | --- | --- | --- | --- | --- | --- |
| ICAM-1 | Mono | Rat | Biolegend | 116102 | 1:100 | O/N | 4℃ |
| CD31 | Poly | Rabbit | Abcam | ab28364 | 1:100 | O/N | 4℃ |
| VCAM-1 | Poly | Rabbit | Abcam | ab134047 | 1:100 | O/N | 4C |
| Mac-2 | Mono | Rat | Cedarlane | CL8942AP | 1:5,000 | O/N | 4C |
| Isolectin-B4, fluorescein | NA | Griffonia Simplicifolia | Vector Laboratories | FL-1201-.5 | 1:100 | O/N | 4C |
| AlexaFluor 488 | Poly | Goat | Invitrogen | A32731 | 1:500 | 1 hour | RT |
| AlexaFluor 546 | Poly | Goat | Invitrogen | A11081 | 1:500 | 1 hour | RT |
| AlexaFluor A647 | Poly | Goat | Invitrogen | A21245 | 1:500 | 1 hour | RT |
| IgG control | Poly | Rat | Santa Cruz | sc-2026 G1319 | Matched by weight | O/N | 4C |
| IgG control | Poly | Rabbit | Vector Laboratories | I-100 | Matched by weight | O/N | 4C |

**Other Reagents – Immunohistochemistry**

| **Reagent** | **Supplier** | **Catalog #** |
| --- | --- | --- |
| Antibody Diluent pH 7.4 | IHC-TEK | IW-1000 |
| TrueVIEW Autofluorescence Quenching Kit | Vector Laboratories | NC1429064 |
| Mounting media: VECTASHIELD with DAPI | Vector Laboratories | 101098-044 |

**Software (all freely available)**

| **Software** | **Use** | **Download** |
| --- | --- | --- |
| ImageJ (FIJI) | Gross lesion analysis | <https://imagej.net/software/fiji/downloads> |
| Olyvia | Olympus file viewer | [https://www.olympus-lifescience.com/en/support/downloads/](https://www.olympus-lifescience.com/en/support/downloads/#!dlOpen=%2Fen%2Fdownloads%2Fdetail-iframe%2F%3F0%5Bdownloads%5D%5Bid%5D%3D847249644) |
| QuPath | Morphometric and immunofluorescence analysis | <https://qupath.github.io/> |
| R | Statistics and data visualization  (packages: dplyr, ggplot2) | <https://cran.r-project.org/bin/windows/base/> |
